# The Impact of Glutathione Administration on Body Weight and Lipid Metabolism in Mice Following Exposure to Mobile Phone Radiation of Frequency 850-1900

**DOI:** 10.1101/2024.05.24.595675

**Authors:** Aliyah Temitayo Ahmed, Abdulhakeem Binhambali, Abdullahi Hussein Umar, Amat Abdoulie Tekanyi, Aisha Ahmad Pate, Amina Nuhu, Tolulope Ipinlaiye, Rabiu AbduSSALAM Magaji

**Affiliations:** Department of Human Physiology, Faculty of Basic Medical Sciences, College of Medical Sciences, Ahmadu Bello University, Zaria, Nigeria; Department of Clinical Science, College of Veterinary Medicine, North Carolina State University, Raleigh, USA; Translation Research in Pain, College of Veterinary Medicine, NCSU, Raleigh, USA

**Keywords:** Mobile phone radiation, Glutathione, Serum lipid profiles, High density lipoprotein, Radiation effect

## Abstract

In an era where electronic advancements surge forward at an unprecedented pace, public apprehension surrounding the health ramifications of radio-frequency (RF) radiation intensifies. This study looked into the impact of chronic exposure to mobile phone radiation, coupled with glutathione supplementation, on serum lipid profiles and body weight in mice—a subject vital for understanding potential health risks in humans and animals respectively.

Thirty-five male mice were divided into seven groups, each subjected to various mobile phone modes and glutathione treatments. Over five weeks, these mice were exposed to 300 missed calls daily, simulating real-world exposure scenarios. Different serum lipid parameters—triglycerides, cholesterol, high-density lipoprotein (HDL), and low-density lipoprotein (LDL)—were monitored and observed throughout the study period.

Remarkably, triglycerides, cholesterol, and LDL levels exhibited no noteworthy statistical variances compared to the control group or among experimental cohorts exposed to diverse range of mobile phone radiation modes or glutathione supplements. However, we noted some little deviations in HDL levels, particularly in the silent and silent+glutathione groups. Moreover, we also noted an unexpected result in weight loss across all groups by week three, which was pronounced by week five, and notably pronounced in the glutathione-administered cohorts. This outcome hints at glutathione’s complex role in mitigating some complications of mobile phone radiation exposure.

In conclusion, the study suggests a potential interplay between glutathione supplementation, mobile phone radiation exposure, and HDL levels in mice—an avenue ripe for further exploration into the understanding of cellular response to modern technological exposures.

## INTRODUCTION

The ubiquitous presence of mobile phones, embraced by approximately 91% of the younger population globally, has revolutionized our lifestyles [1]. These devices have seamlessly integrated into daily routines, becoming indispensable tools for communication and information access. However, as mobile phone usage continues to soar, concerns about their potential impact on human health have intensified [2, 3]. With mobile phones pervading every corner of the globe, exposure to electromagnetic field radiation (EMR) has become a constant presence in modern society [4].

Radiations consist of both electric and magnetic fields, originating from natural and man-made sources. EMR is present in various scenarios of everyone’s life, with some of the most common sources being solar radiation, household electric currents (e.g., mobile phones, television sets, Wi-Fi, microwave ovens, computers), and telecommunication antennas [5–11].

Electromagnetic frequency (EMF) has numerous biochemical consequences, including the degradation of large cell molecules and imbalances in physiological ionic equilibrium. EMF exposure is associated with increased formation of reactive oxygen species (ROS) [12], which can cause injury to cellular constituents such as lipids, proteins, and DNA. It has been shown that electromagnetic radiation influences living organisms by producing or increasing ROS, leading to various physiological and biochemical effects, including DNA damage [13]. Analog telephones typically use frequencies between 800 and 900 MHz, while digital telephones use frequencies between 1850 and 1990 MHz, and microwave ovens operate at a frequency of 2450 MHz. The 900 MHz EMF emitted by mobile phones is commonly used in low-income countries.

However, cell phone radiation, a form of electromagnetic radiation known as microwaves, comprises electric and magnetic waves that travel through space at the speed of light [14]. Concerns have been raised by researchers regarding the release of EMF from mobile phones into the atmosphere, suggesting potential harmful effects on human health [15, 16]. Indeed, the adverse impacts of EMF have become a pressing global issue, with reports of detrimental effects on various organs such as the brain, ear, and reproductive organs [17, 18, 5, 19, 20].In today’s digital era, the proliferation of cell phone usage is on the rise, contributing to a growing addiction among individuals [21]. Over the past two decades, mobile phone subscriptions have surged from 12.4 million to over 5.6 billion globally, reaching approximately 70% of the world’s population [74]. However, alongside this widespread adoption, concerns regarding its impact on public health have emerged. There are numerous reports of both mental and physical health hazards associated with cell phone use across all age groups, with some effects proving to be severe, including the heightened risk of cancers [22].

Electromagnetic Radiation (EMR) is a dynamic force of energy propagation, characterized by photons exhibiting both particle and wave properties, traveling at the speed of light. These waves bear energy and transmit it upon interaction with matter. The energy content of EMR is directly linked to its frequency and inversely related to its wavelength, meaning that waves with shorter wavelengths possess higher energy levels. EMR encompasses a spectrum ranging from radio waves to microwaves, infrared, visible light, UV, and radiographs, each with varying energy levels [23]. Both EMR and particulate radiation have the ability to generate ion pairs through their interaction with matter. Ionizing radiation, including alpha particles, neutrons, beta rays, charged nuclei, and positron radiation, aids in producing high-quality images for medical diagnostics and is utilized in cancer treatment as well [24].

The impact of EMF extends to biochemical processes, including the degradation of large cellular molecules and disruptions in ionic equilibrium. Exposure to EMF has been linked to heightened formation of reactive oxygen species (ROS) [12] as stated earlier, ROS generation can also occur aside from the previously mentioned EMF, including, radioactivity, stress, cigarette smoke, and redox processes within the body [5]. It is also important to know that the adverse effects of mobile phone radiation exposure on vital organs like the kidney and testis remain poorly understood, as does its impact on serum lipids, which play crucial roles in diseases such as cardiovascular disorders [13]. While interest in evaluating the health effects of non-ionizing radiation, particularly radiofrequency (RF) electromagnetic fields, has grown significantly in recent decades, fueled by the widespread use of mobile telecommunications worldwide, the exact mechanisms and consequences of such exposure are still being unraveled. Mobile networks operate on varying frequencies across different regions, with Global System for Mobile Communications (GSM) networks typically utilizing the 900 and 1800 MHz bands [30].

Among the potential effects of radiofrequency radiation on the human body, research has predominantly focused on its impact on the brain, cancer incidence, and fertility. This includes investigations into enzyme induction, neurological symptoms, toxicological effects, genotoxicity, carcinogenicity, and alterations in sperm cell fertility. The literature extensively examines the influence of radiofrequency radiation on various cellular processes, including mitochondrial function, apoptosis pathways, heat shock proteins, free radical metabolism, cell proliferation, differentiation, DNA damage, and plasma membrane integrity [31–36]. Despite these efforts, gaps remain in our understanding of the full spectrum of health effects associated with radiofrequency radiation exposure, necessitating further research to elucidate its potential risks and inform regulatory measures to protect public health.On the other hand, non-ionizing radiation, though lacking in ionizing power, can still pose risks with prolonged exposure, a crucial factor when considering its interaction with the human body [26]. The consensus among many researchers is that non-ionizing radiation carries significant hazards for public health [27].Termed electromagnetic pollution, the potential health impacts of EMF exposure are increasingly recognized [28]. Non-ionizing radiation spans from extremely low frequency (ELF) to visible light, while ionizing radiation ranges from higher frequencies beyond visible light[29].The proliferation of electric EMF from modern technologies, including everyday appliances and mobile devices, poses a tangible risk to human well-being as explained earlier.

Glutathione, on the other hand, is a vital tripeptide composed of cysteine, glycine, and glutamic acid, serving as a cornerstone in numerous bodily functions, from combating oxidative stress to bolstering immune responses [37]. Its significance extends to mitigating toxin accumulation by aiding in the detoxification process, particularly in the liver’s conversion and elimination of harmful compounds like mercury and persistent organic pollutants (POPs). Functioning as a potent antioxidant, glutathione is ubiquitous across various organisms and cell types, primarily existing in either its reduced (GSH) or oxidized (GSSG) form [38, 39]. Under normal circumstances, the majority of glutathione remains in its reduced state, pivotal for maintaining cellular health and redox balance. Studies by Markov *et al.* [40], have explored the impact of radio frequency electromagnetic waves (RF-EMW) on human health, noting a potential correlation between long-term exposure and reduced body weight. While RF radiation, falling within the spectrum of 0.5 MHz to 100 GHz, may affect certain organs and systems, the precise health risks from various sources of non-ionizing radiation, including those emitted by mobile phones, remain less understood [41].Research by NRC and Roberts *et al.* [42] has indicated that exposure to RF/MW radiation can influence several hematological variables, both in animal models and humans. This underscores the need for further investigation into the potential health implications of non-ionizing radiation exposure, particularly in light of its pervasiveness in modern society.

In animal research, the welfare of experimental subjects is of paramount importance. Criteria such as a 20 percent reduction in body weight are often designated as indicators of severe suffering, guiding humane endpoint decisions [43]. Previous studies have underscored the potential impact of low electric fields on mice, revealing diminished body weight and reduced water intake following one month of exposure to harsh conditions [44].

Mobile phone users in various countries and continents are exposed to different frequencies. EMF exposure depends on the frequency of the cell phone [45]. Furthermore, there are indications that mobile phone radiation may also influence body weight in experimental animals. In our contemporary era, where mobile phone usage is common and its potential health implications are a subject of concern, we endeavor to explore the effect of such radiation on serum lipid and the interplay between glutathione and body mass in mice subjected to chronic mobile phone radiation exposure. Through this investigation, we aim to shed light on the complex relationship between glutathione, mobile phone radiation, and serum lipid in experimental models. We evaluated the effects of EMF on the serum lipid profiles and the body weight and the potentially harmful effects of EMF on these parameters of mice model.

## MATERIALS AND METHOD METHODOLOGY

### Animals and grouping

The animals involved in the study were maintained and used in accordance with the Animal Welfare Act and the Guide for the Care and Use of Laboratory animals prepared by the Ahmadu Bello University Committee on Animal Use and Care (ABUCAUC). And ethical approval was obtained from the ABUCAUC.

At the start of the experiment, thirty five (35) apparently healthy adult male Mice between the ages of 8-12 weeks were obtained from the Department of Human Physiology, College of Basic Medical Sciences, Ahmadu Bello University, Zaria, Nigeria animal breeding section. They were housed in standard polypropylene cages and were maintained and allowed free access to feed and water 24/7 and 12h alternate day light and darkness. The animals were allowed to acclimatize to the environment of the behavioral laboratory for two weeks before the commencement of the experiment and they were provided with feeds and water adlibitum.

The animals were randomly selected and divided into seven (7) groups of five (n=5) mice per group as follows: Group I: Normal control group, Group II: Group exposed to mobile phone radiation in ringtone mode, Group III: Group exposed to mobile phone radiation in vibration mode, Group IV: Group exposed to mobile phone radiation in silent mode, Group V: Group exposed to mobile phone radiation in ringtone mode plus glutathione, Group VI: Group exposed to mobile phone radiation in vibration mode plus glutathione and the last group, Group VII were exposed to mobile phone radiation in silent mode plus glutathione.

The animals in groups II to VII were all exposed to 4hour mobile phone radiation of 900 hertz and an average of 300 missed calls per day for 5 weeks. The group I as the normal control is not exposed to any form of radiation or glutathione.

### Radiation exposure

Using Itel mobile phone (it2160) with specific absorption rate (SAR) of 2W/kg and 2G Network Bands, of frequency 850-1900 MHz, the animals in groups II to VII were all exposed to 4 hours mobile phone radiation of an average of 300 missed calls per day for six (6) weeks. Animals in groups V, VI and VII were administered glutathione (250 mg/kg) thirty minutes prior to mobile phone radiation daily.

### Determination of body weight

The body weight is taken early in the morning (between 7am-9am) prior to their feeding. Animals were individually weighed on a weekly basis (Wednesdays) in order to detect any changes in their body weights. Before administration of glutathione and exposure to radiation, the animals were weighed with a sensitive weighing balance (Uline Balance Scale – H-9886, 7.5X5.7 square, 2,200 g x 0.1 g) Glutathione at a dose of 250 mg/Kg, was administered orally to each Mice daily to group V-VII before exposure to radiation for 6 weeks. Animals in group II - IV were exposed to mobile phone radiation only, for 4 hours daily throughout the six weeks radiation without Glutathione

### Determination of lipoproteins

#### Triglyceride

0.5ml of working reagent was added to a set of clean labeled test tubes and two of the test tubes were labeled standard 1 and 2 while another was labeled “blank”.0.1ml of the sample was added to the approximately labeled test tube .mixed and incubated for 30 minutes at 37°, the absorbance of the sample and standard were read against the reagent blank at the wavelength of 546nm.

#### Total Cholesterol (TC)

0.5ml of working reagent was added to a set of clean labeled test tubes and two of the test tubes were labeled standard 1 and 2 while another was labeled “blank”.0.1ml of homogenate was added to the approximately labeled test tube .mixed and incubated for 30 minutes at 37°, the absorbance of the sample and standard were read against the reagent blank at the wavelength of 546nm.

#### High-density lipoproteins (HDL)

0.1 ml of homogenate sample was mixed with 0.5ml of working reagent (R1), it was mixed and incubated for exactly 30 minutes for 37°, 0.5ml of the second reagent (R2) was mixed and allowed to stand for 20 minutes at 20-25°C, then 5.0ml of sodium hydroxide was added and mixed, the absorbance of the sample (A sample) against the reagent blank was read after 5 minutes at the wavelength of 546nm 21

#### Low-density lipoproteins (LDL)

LDL cholesterol was calculated from measured values of total cholesterol, triglycerides and HDL-cholesterol according to the relationship:

[LDL-chol]= [total chol] – [HDL-chol] – [TG]/5

Where [TG]/5 is an estimate of VLDL-cholesterol and all values are expressed in mg/dl

## RESULTS

### WEEKLY WEIGHT PROFILES

In the first week, the impact of mobile phone radiation and glutathione on mean body weight in mice is illustrated in [Figure 1]. The number of open entries recorded were as follows: control (19.2400 ± 1.00080), ringtone (17.8600 ± .56798), vibration (18.8200 ± .55624), silent (16.7000 ± .92304), ringtone + glutathione (20.9200 ± .1.01114), vibration + glutathione (19.3000 ± 1.31567), and silent + glutathione modes (20.9000 ± 1.52381). No statistically significant difference (p > 0.05) was observed when comparing the number of entries in the control group to all other groups. The impact of mobile phone radiation and glutathione on body weight in mice, as depicted in [Figure 1], was also examined. The number of open entries recorded were: control (21.0400 ± .66076), ringtone (18.4800 ± .65985), vibration (19.8400 ± .63135), silent (17.3200 ± .49739), ringtone + glutathione (21.3000 ± .86487), vibration + glutathione (20.5600± 1.10526), and silent + glutathione modes (20.5800 ± 1.33768). There was no statistically significant difference (p > 0.05) in the number of entries between the control group and all other groups, except for the silent, ringtone+GSH, and silent=GSH. Similarly, the effects of mobile phone radiation and glutathione on body weight in mice are displayed in [Figure 1]. The number of open entries were: control (24.5600 ± .46648), ringtone (22.6200 ± .40423), vibration (22.1800 ± .69957), silent (21.8200 ± .45869), ringtone + glutathione (20.3600 ± .86487), vibration + glutathione (19.4200 ± 1.15039), and silent + glutathione modes (22.9000 ± 1.10770). There was no statistically significant difference (p > 0.05) in the number of entries between the control group and all other groups, except for ringtone + GSH, vibration + GSH, and silent+GSH. Furthermore, the impact of mobile phone radiation and glutathione on body weight in mice is showcased in [Figure 1]. The number of open entries recorded were: control (26.1000 ± .90277), ringtone (22.1600 ± .59380), vibration (22.5200 ± .91291), silent (21.7400 ± .58703), ringtone + glutathione (23.3800 ± .75525), vibration + glutathione (22.0340 ± 1.14471), and silent + glutathione modes (20.5200 ± 1.26270). A statistically significant difference (p<0.05, ANOVA) was observed when comparing the number of entries in the control group to all other groups, except for vibration and ringtone+GSH. Lastly, the impact of mobile phone radiation and glutathione on body weight in mice, illustrated in [Figure 1], indicates the following number of open entries: control (27.2200 ± .34264), ringtone (22.8600 ± .63372), vibration (25.1200 ± .58856), silent (23.4000 ± .70071), ringtone + glutathione (23.2400± .83403), vibration + glutathione (22.3400± 1.01371), and silent + glutathione modes (21.3600 ± 1.31095). A statistically significant difference (p<0.05, ANOVA) was noted when comparing the number of entries in the control group to all other groups.

**Figure 1:**
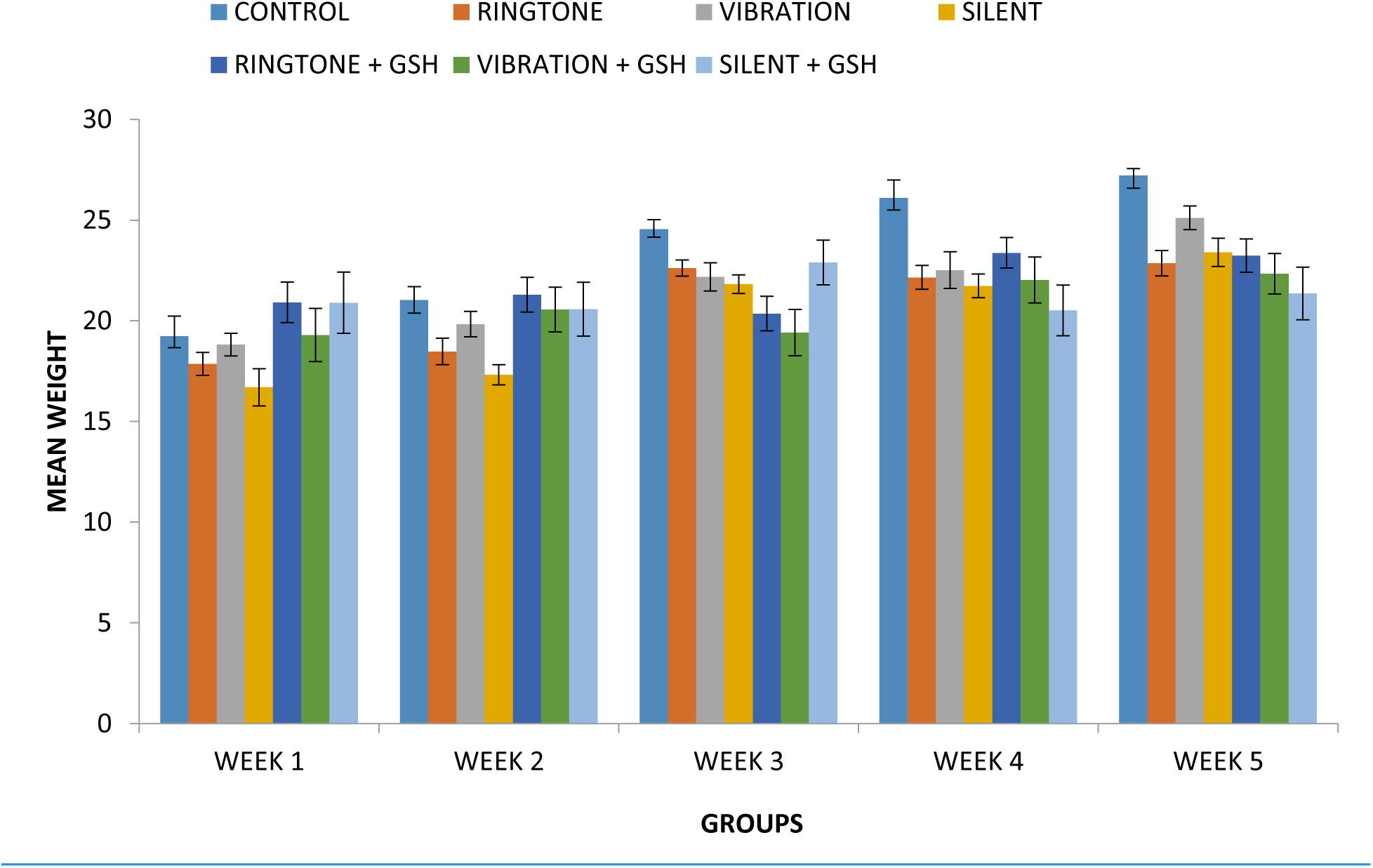
Effects of mobile phone radiation and glutathione on body weight in mice

## SERUM LIPID PROFILES

### Triglycerides

The effect of mobile phone radiation and glutathione on triglycerides level in mice is shown in [Figure 2]. The concentration level were; control (109.5 ± 18.9), ringtone (110.4 ± 6.8), vibration (111.7 ± 7.2), silent (85.5 ± 5.3), ringtone + glutathione (115.6 ± 10.3), vibration + glutathione (108.5 ± 6.0) and silent +glutathione modes (81.5 ± 6.0). There was no statistical significance (*p* > 0.05) difference when the concentration level of control group was compared to all the groups.

**Figure 2:**
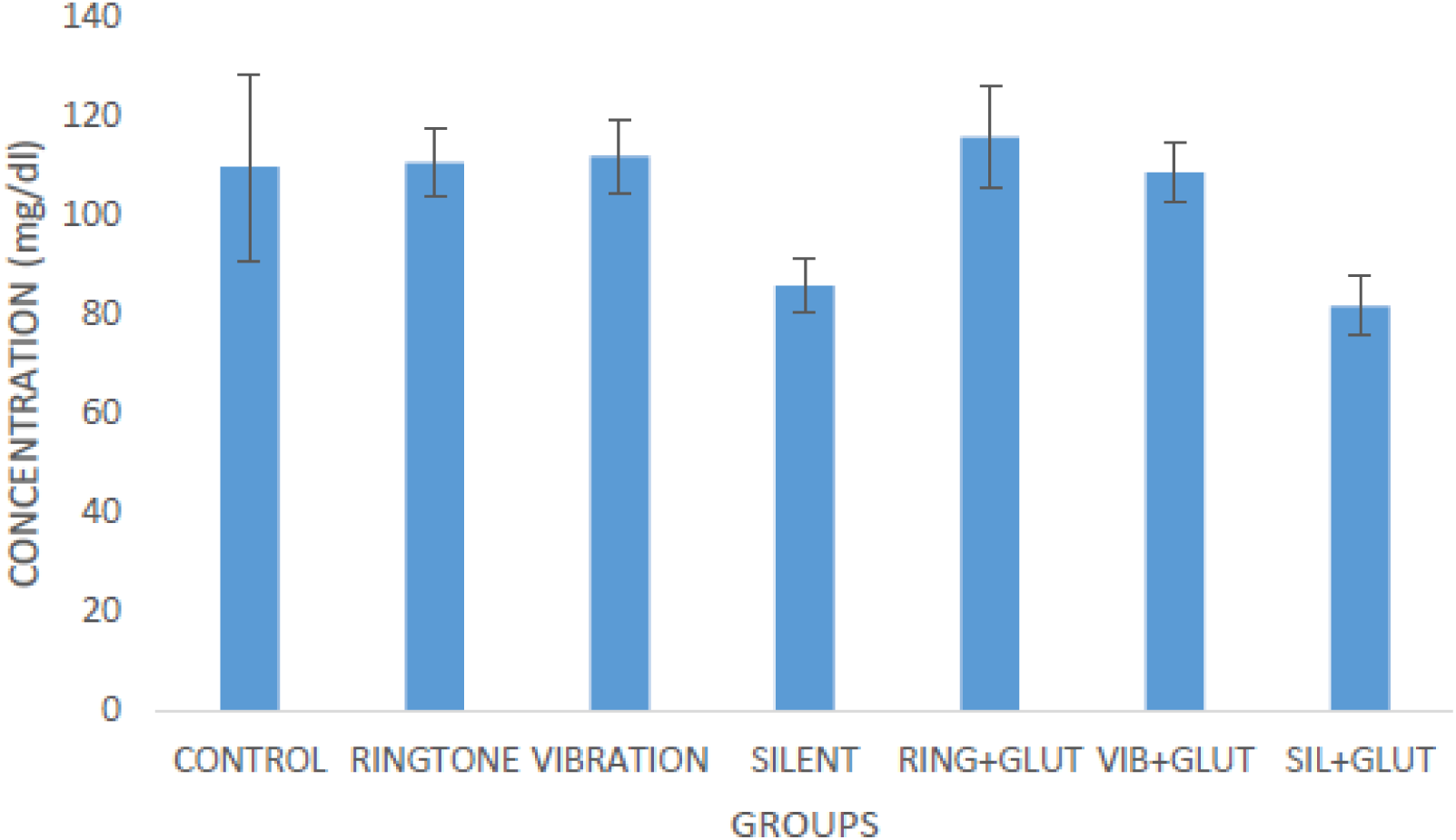
Effects of mobile phone radiation and glutathione on triglycerides level in mice. (Mean ± S.E.M, n = 7). Glut=Glutathione, Sil= Silent, Vib=Vibration, Ring=Ringtone There were no statistical significance differences between the groups (*P* > 0.05 ANOVA).

### Cholesterol

The effects of mobile phone radiation and glutathione on cholesterol level in mice are shown in [Figure 3]. The concentration level were; control (147.4 ± 14.8), ringtone (122.7 ± 2.5), vibration (166.0 ± 9.3), silent (140.3 ± 13.0), ringtone + glutathione (115.8 ± 20.5), vibration + glutathione (134.3 ± 14.4) and silent +glutathione modes (123.7 ± 17.1). There was no statistical significance (*p* > 0.05) difference when the concentration level of control group was compared to all the groups.

**Figure 3:**
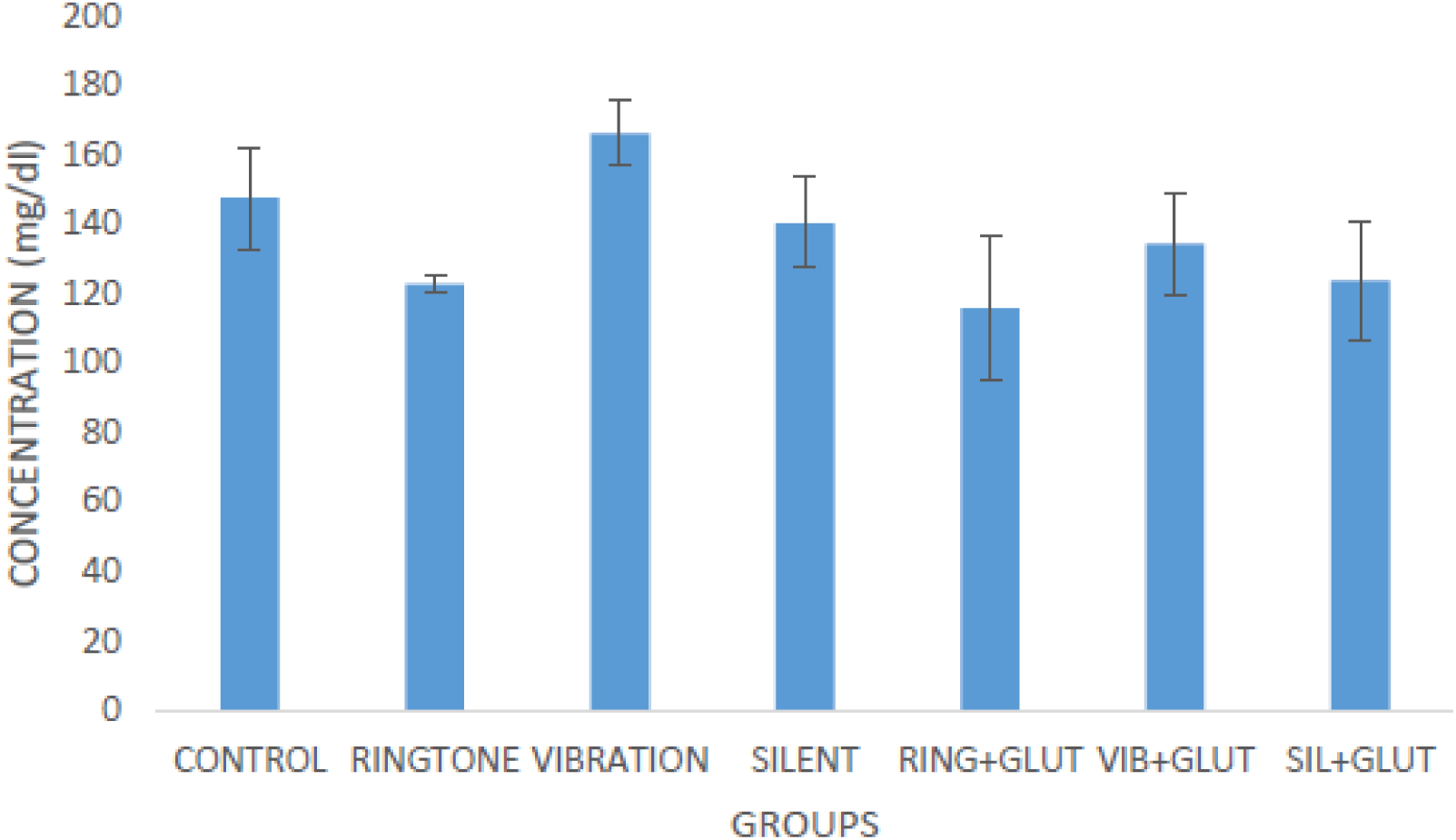
Effects of mobile phone radiation and glutathione on cholesterol level in mice. (Mean ± S.E.M, n = 7). Glut=Glutathione, Sil= Silent, Vib=Vibration, Ring=Ringtone There were no statistical significance differences between the groups (*P* > 0.05 ANOVA).

### High-density lipoproteins

The effects of mobile phone radiation and glutathione on high-density lipoproteins (HDL) level in Mice are shown in [Figure 4]. The concentration level were; control (111.2 ± 5.5), ringtone (79.3 ± 3.4), vibration (108.8 ± 6.2), silent (116.5 ± 7.4), ringtone + glutathione (117.0 ± 8.7), vibration + glutathione (124.8 ± 6.2) and silent +glutathione modes (86.9 ± 4.5). There was statistical significance (*p* < 0.05) difference when the concentration level of control group was compared to ringtone and silent+glutathione groups and no significant differences was seen in the remaining groups.

**Figure 4:**
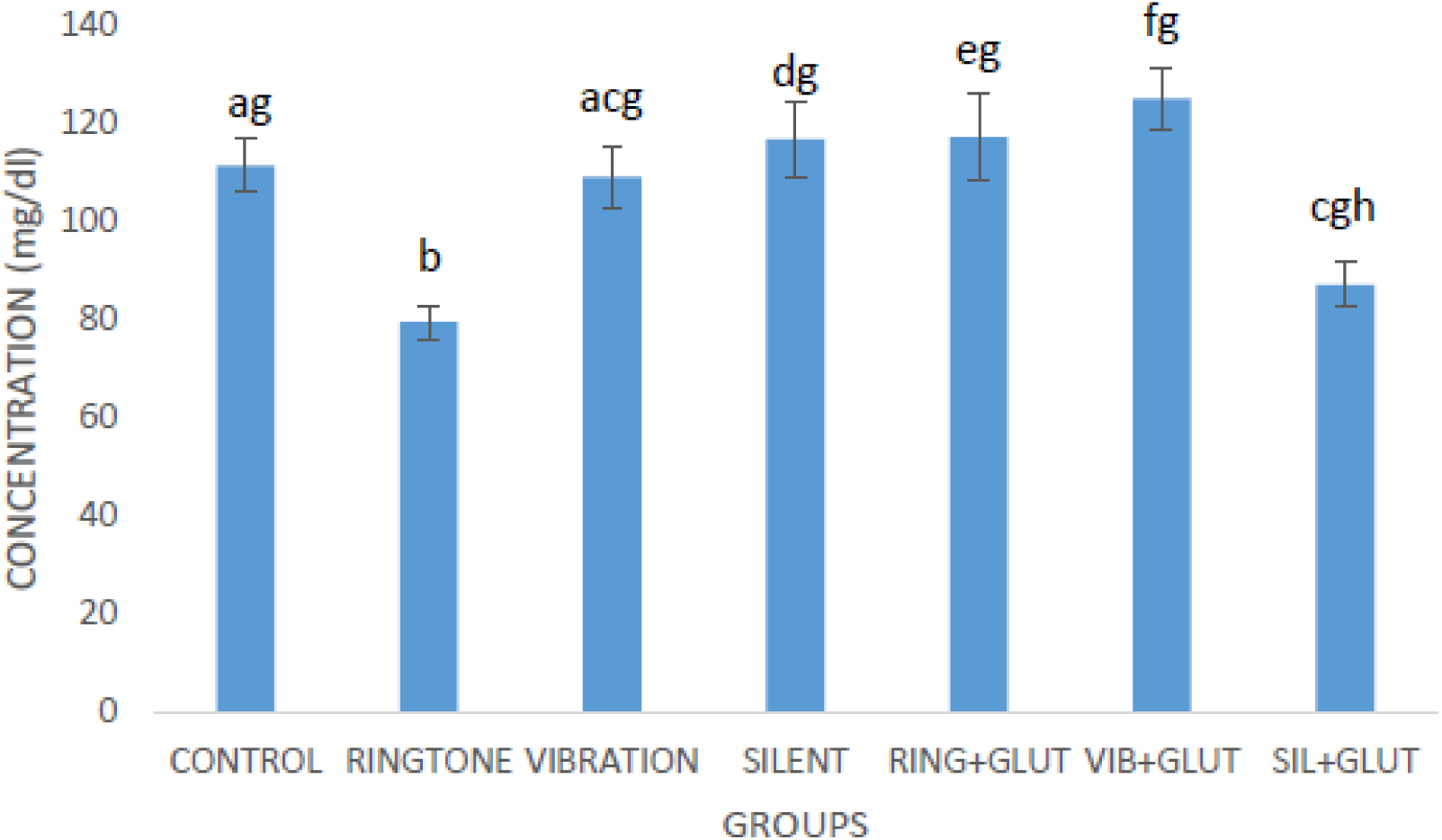
Effects of mobile phone radiation and glutathione on high-density lipoproteins level in mice. Bars with different letters are significant (Mean ± S.E.M, n = 7). Glut=Glutathione, Sil= Silent, Vib=Vibration, Ring=Ringtone There was statistical significance differences between the groups (*P* <0.05 ANOVA).

### Low-density Lipoprotein

The effects of mobile phone radiation and glutathione on low-density lipoproteins (LDL) level in mice are shown in [Figure 5]. The concentration level were; control (65.2 ± 16.7), ringtone (23.3 ± 2.9), vibration (46.8 ± 6.5), silent (28.7 ± 3.0), ringtone + glutathione (23.0 ± 7.4), vibration + glutathione (35.1 ± 10.3) and silent +glutathione modes (32.7 ± 16.4). There was no statistical significance (*p* > 0.05) difference when the concentration level of control group was compared to all the groups.

**Figure 5:**
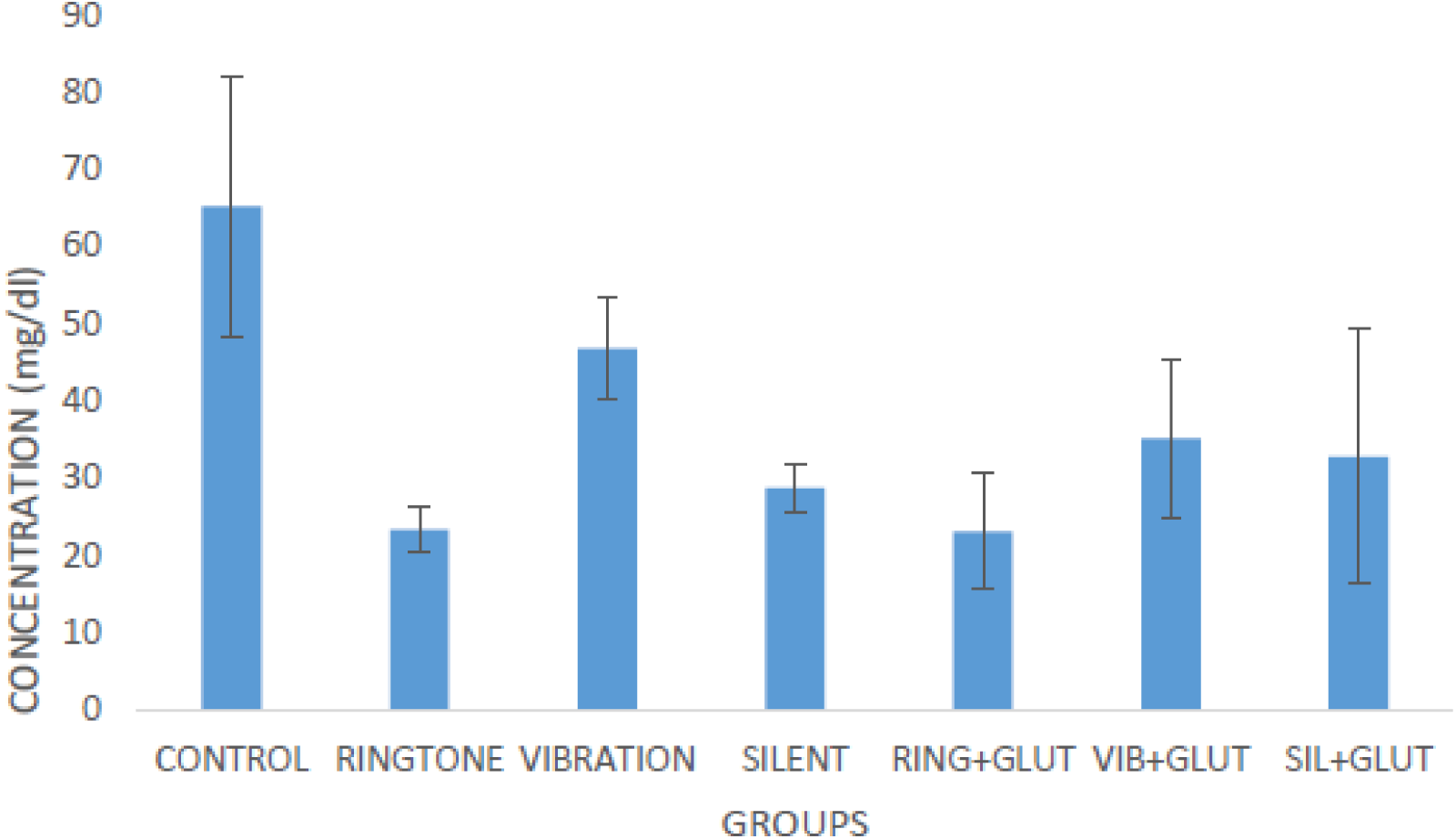
The effects of mobile phone radiation and glutathione on low-density lipoproteins level in mice. (Mean ± S.E.M, n = 7). Glut=Glutathione, Sil= Silent, Vib=Vibration, Ring=Ringtone There were no statistically significance differences between the groups (*P* > 0.05 ANOVA)

## DISCUSSION

Many studies conducted in recent years have reported various biological effects, spanning epidemiological and clinical investigations in humans, as well as experimental research involving rodents, flies, and cell cultures exposed to phone radiation[46]. However, despite this extensive research, little attention has been paid to serum lipid parameters and body weight reduction to radiation exposure. In today’s era, mobile phones have become indispensable tools in our daily lives, serving critical roles in business, commerce, education, and emergency communication. Nevertheless, the proliferation of EMR from sources such as cell phone towers, TV, and FM towers has raised concerns about potential health hazards [47]. Studies by Hocking *et al.* in Australia have linked the proximity of individuals to such towers with higher incidences of leukemia, tumors, and cancer, highlighting the need for further investigation into these health risks [7, 48].

Lipid peroxidation, a consequence of oxidative stress induced by radiation, entails the degradation of lipids when reactive oxygen species (ROS) generated by radiation interact with polyunsaturated fatty acids in the phospholipid membrane bilayer [49]. Glutathione, a vital cellular antioxidant, plays a crucial role in maintaining cellular health. Notably, it has been observed to significantly impact high-density lipoprotein levels [49]. Found abundantly in foods like onions, garlic, avocados, and cruciferous vegetables, glutathione and its precursors, such as l-cysteine and lipoic acid, function as potent scavengers of free radicals, thereby protecting cells from oxidative damage. Moreover, glutathione aids in preserving proteolytic enzymes in the cortical lens fibers, with studies dating back to the late 1960s highlighting its diminished levels in mature cataracts, suggesting its involvement in cataract formation [50].

As a critical antioxidant in various organisms, glutathione prevents damage to cellular components caused by ROS, including free radicals and peroxides. Its sulfur compounds confer a lipid-lowering effect, with lower levels of glutathione associated with an increased risk of cardiovascular disease [51]. Consequently, supplementation with glutathione precursors or engaging in regular exercise to stimulate glutathione production may confer cardiovascular benefits during different pathophysiological conditions [52].

While numerous studies have demonstrated the antioxidant effects of glutathione in various metabolic contexts, its role in mitigating oxidative stress induced by electromagnetic field (EMF) exposure remains unexplored in the literature. In this study, we investigated the effects of 900- MHz radiofrequency EMF exposure on oxidative stress in adult mice by assessing parameters such as triglycerides, cholesterol, HDL, LDL and body weight.

Numerous studies employing robust methodologies have underscored the potential of electromagnetic fields (EMF) to induce adverse health effects. In the present study, we reported the repercussions of exposure to mobile phone radiation, specifically exploring the impact of different modes (ringtone, vibration, silent, etc.) on lipid profiles and weight in mice. We investigated potential interactions between radiation and its effects on various groups of mice, aiming to discern any discernible patterns or differences. Our findings unveiled new insights into the relationship between mobile phone radiation and lipid parameters. While triglycerides (TGCL), cholesterol (CHL), and low-density lipoprotein (LDL) levels did not exhibit statistical differences compared to the control group, high-density lipoprotein (HDL) levels showed a significant difference [Figure 3]. Additionally, we scrutinized the effect of radiation on body weight, observing notable changes across all groups at week 5, with variations also evident at weeks 3 and 4 [Figures 1]. These observations align with previous studies by Wilson *et al*.[54] and Sokolovic *et al*.[55], highlighting the short-term effects of electromagnetic fields on body mass [54, 55, 56].

Findings reported by Borek *et al*. [65], noted that when rats are exposed to 900-MHz EMF, they exhibited increased protein oxidation and lipid peroxidation in the brain. However, administration of 500 mg/kg/day dissolved sulfur compound (garlic) mitigated these effects, bringing the levels back to normal. In our investigation using glutathione, which is also found in garlic(as reported in their study), we found that glutathione supplementation did not attenuate the oxidative effects of EMF exposure on body weight at any point in time; rather the group with glutathione showed more weight loss from week two and more pronounced at week five than the control group and the groups that were not given glutathione. This was a surprising thing to us, with the known fact that glutathione reduces oxidative stress, but in our case the result was contradictory.

Also interesting to know, our study’s duration and methodology yielded results consistent with the notion that electromagnetic field exposure may induce short-term fluctuations in body weight, as observed in similar experiments with rats [56]. Also noted, our findings was similar with the report by Shabat *et al*[57], which noted a decrease in initial mean weight in groups exposed to high electromagnetic radiation [57]. Likewise the findings from the studies by Hassan *et al*. [13] and Lee *et al*. [58], which found no significant weight differences in rats exposed to magnetic fields [13, 58] was contradicting to our findings which noticed a significant weight loss in some groups from week 3 and more pronounced at week five and even more especially from the groups that were given glutathione. Others have also reported an increase in body weight with long-term exposure to electromagnetic radiation [59] and our finding does not agree with this point. The reduction in body weight post-EMF exposure may be attributed to direct damage and excessive production of reactive oxygen species (ROS) induced by electromagnetic radiation, which glutathione may not adequately scavenge effectively during the period of radiation exposure [60].

Our study also shed light on the significance of HDL-associated proteins, highlighting the interplay between serum lipid levels and tumor genesis [64]. While our findings regarding serum HDL-c levels contradicted those of Wang *et al*.[62], who reported no statistical significance, although it’s essential to note that their focus was primarily on breast cancer, which differs from our study’s scope [62].Moreover, our study did not find statistical significance in cholesterol and triglyceride levels, aligning with findings from other studies suggesting that these parameters may not be directly associated with disease initiation or progression from radiation exposure [63, 64, 65].

In summary, our findings revealed that exposure to a 900-MHz EMF led to increased serum HDL in mice and prompted a loss of body weight by week 3. Glutathione administration mitigated EMF-induced weight loss in mice, particularly evident at week 5. However, we advocate for further research, particularly focusing on these parameters with further studies on liver enzymes and serum electrolytes, to deepen our understanding of the complexities surrounding EMF exposure and its potential health implications.

## ACKNOWLEDGEMENT

This case was carried out without direct funding, and the authors would like to thank the Head and the entire staffs of the Department of Human Physiology, College of Medical Sciences, Ahmadu Bello University, Zaria, for their support and provision of necessary facilities. The authors also acknowledge the indispensable contributions of the Technicians at the Department, whose dedication greatly contributed to the success of the study.

## AUTHOR’S CONTRIBUTIONS

All authors contributed greatly to the success of this work

